# Mesophotic Coral Ecosystems Inside and Outside a Caribbean Marine Protected Area

**DOI:** 10.1101/241562

**Authors:** Erika Gress, Maria J Arroyo-Gerez, Georgina Wright, Dominic A Andradi-Brown

## Abstract

Recent widespread shallow coral reef loss has led to calls for more holistic approaches to coral reef management, requiring inclusion of all ecosystems interacting with coral reefs in management plans. Yet almost all current reef management is biased towards shallow reefs, and overlooks that many reef species can also be found on mesophotic coral ecosystems (MCEs; reefs 30 −150 m). This study presents the first detailed quantitative characterisation of MCEs off Cozumel, in the Mexican Caribbean and provides insights into their general state. We investigate whether MCEs within the marine park have similar ecological communities to mesophotic reefs outside protection, despite widely recognised shallow reef impacts outside the protected area. Results show some taxon specific differences in MCE benthic communities between sites within the protected area and areas outside; although overall communities are similar. Regardless of protection and location, and in contrast to shallow reefs, all observed Cozumel MCEs were continuous reefs dominated by calcareous macroalgae, sponges, octocorals, and black corals. Hard corals were present on MCEs, but at low abundance. We found that 42.5 % of fish species recorded on Cozumel could be found on both shallow reefs and MCEs, including many commercially-important fish species. This suggest that MCEs may play a role in supporting fish populations. However, regardless of protection status and depth we found that large-body fishes (>500 mm) were nearly absent at all studied sites. MCEs should be incorporated into the existing shallow-reef focused management plan in Cozumel, with well informed and implemented fisheries and harvesting regulations.

## Introduction

Coral reef ecosystems border nearly a sixth of global coastlines [1], contain thousands of species [2], and play a crucial food security role for millions of people [3]. Goods and services from coral reefs have been estimated to be worth >3 billion USD annually [4]. Yet shallow coral reefs face widespread threats, both from local scale impacts (e.g. over-fishing, pollution) and from large scale impacts (e.g. coral bleaching, ocean acidification) [3,5–7]. In the face of such threats, many recent conservation efforts have focused on maintaining shallow reef resilience [8,9] combining the ability of reefs to both resist stressors and recover from damage following impact [9,10]. Yet, little consideration has been given to the role of deeper reef refuge habitats [11]. eeper light-dependent coral ecosystems, known as mesophotic coral ecosystems (MCEs), are found from approximately 30-150 m and are known to have high species diversity [12,13] including scleractinian corals, sponges, octocorals, black corals, and macroalgal species [6]. It has been suggested that MCEs may be less exposed to anthropogenic impacts than adjacent shallow reefs [6,11].

Ecological research on MCEs has increased recently, but MCEs remain under-studied because of technical, logistical and financial challenges associated with accessing them [7,9,11,14–16]. Studies show that upper-MCEs (30-60 m) often contain species found on shallow reefs [7,11,15,17–19], while lower-MCEs (60-150 m) may contain more deeper-water specialist species [11]. The ‘deep reef refugia hypothesis’ (DRRH) suggests that MCEs are protected from disturbances that affect shallow reef areas, such as rising water temperatures and coastal development [17,20]. In addition, mesophotic reefs are, in some cases, protected from direct fisheries exploitation [21,22], with larger individual fish recorded at near-MCE depths [21,23]. Despite this, MCEs face many similar threats to shallow reefs [24], with examples of overexploitation from targeting economically important fishes [21,25,26] and black corals [27–29] on MCEs. In addition, other processes such as sedimentation because of adjacent human development can lead to MCE habitat degradation [15].

There has been an increase in discussion about the relevance of MCEs [30,31] and their role in reef resilience and conservation [17]. However, the few examples of MCE management are focused on small areas and/or single taxa. Black corals (Antipatharians) for use in the jewellery trade [32] has led to specific harvesting regulations in Hawaii, for example [33]. Black corals are long-lived, ahermatypic corals, that are crucial habitat-forming species on some MCEs because of their complex structure and their ability to form dense beds which other fish and invertebrate species associate with [27,28,33,34]. In response to this, Antipatharians have been regulated by CITES Appendix II since 1981 [35].

There are even fewer examples of MCEs being integrated into broader reef management. A recent exception is the Coral Sea Reserve in Eilat (Gulf of Aqaba, Red Sea) where following MCE documentation, an existing marine park boundary was moved to 500 m further offshore, and to 50 m depth to incorporate MCEs into the protected area [9,17]. Other MCE areas, such as the *Oculina* reefs off the Florida coast have received direct protection through establishment of a new marine protected area after surveys indicated the damage caused by trawling in the area [24,26,36]. Even with very limited MCE data, is possible to integrate MCEs into marine protected areas. For example, on the Great Barrier Reef, MCEs became incorporated within the management plan by ensuring representation of different geological seabed features when conducting park zonation [37]. These approaches fit with the holistic view of reef management recently advocated [9,17].

In this study, we assess shallow reef and MCE benthic and fish communities within the Cozumel National Marine Park and adjacent areas with no protection near to the main tourism development. Shallow reefs are reported to be more degraded in the area without protection [38,39]. We investigate whether MCEs within the marine protected area (MPA) retain similar ecological communities to MCEs outside the MPA. These data will help to serve as a baseline for future studies and also provide insight into the role of these deep reefs as refuges.

## Methods

### Study site

Surveys were conducted around Cozumel, Mexico, an island located 16.5 km off the east coast of the Yucatan peninsula at the northern extent of the Mesoamerican Reef (Figure 1). There are extensive fringing coral reef ecosystems off the west coast of Cozumel, that are well recognized for their biological and socioeconomic importance [40,41]. They are heavily visited by recreational SCUBA divers, with reef related tourism contributing significantly to the island and the whole region’s economy. In 2015, the port of Cozumel received 3.8 million passengers that arrived on 1,240 vessels – more than anywhere else in the world [42]. The reefs of Cozumel are under two protection regimes: a National Marine Park in the southwest, and the Flora and Fauna Protected Area in the north and east coasts (Figure 1). The National Marine Park was decreed in 1996 and is 11,987 ha in area; it is zoned to allow only recreational SCUBA diving and other tourism (including sport fishing) in intensive use areas containing shallow coral reefs, while hook and line fishing is allowed in other less intensive use areas [43]. Cozumel reefs are also now part of the most recently decreed protected area along the Mexican Caribbean that includes approximately 57,000 km^2^ of marine habitats [42]. The Healthy Reefs Initiative (HRI), an international organization that monitors the Mesoamerican Reef has classified the shallow reefs of Cozumel contained within the National Marine Park as in ‘very good’ condition [44]. For Cozumel, their data shows hard coral (scleractinian) coverage at 20-40 %, and the presence of economically important species such as large groupers and snappers [44]. The Flora and Fauna protection, designated in 2012, covers the east and north coasts of the island and it has a different protection regime with only a core zone of 470 ha that is fully no-take for fisheries [45]. The majority of Cozumel reefs are contained within one of these two protection schemes, with the only area of reef without any protected status adjacent to the main development on the island (Figure 1). Here the development of vessel terminals and tourism infrastructure adjacent to the reef is known to have caused widespread shallow reef degradation [38,39,46], including declines in hard coral cover from 44 % to 4 % over the period 1995-2005 [38]. Cozumel is renown for the black coral jewellery industry since the early 1960’s. Antipatharian beds were widely found at upper-mesophotic depths (30-60 m), but were not properly documented prior to overexploitation [47–49].

**Figure 1.**
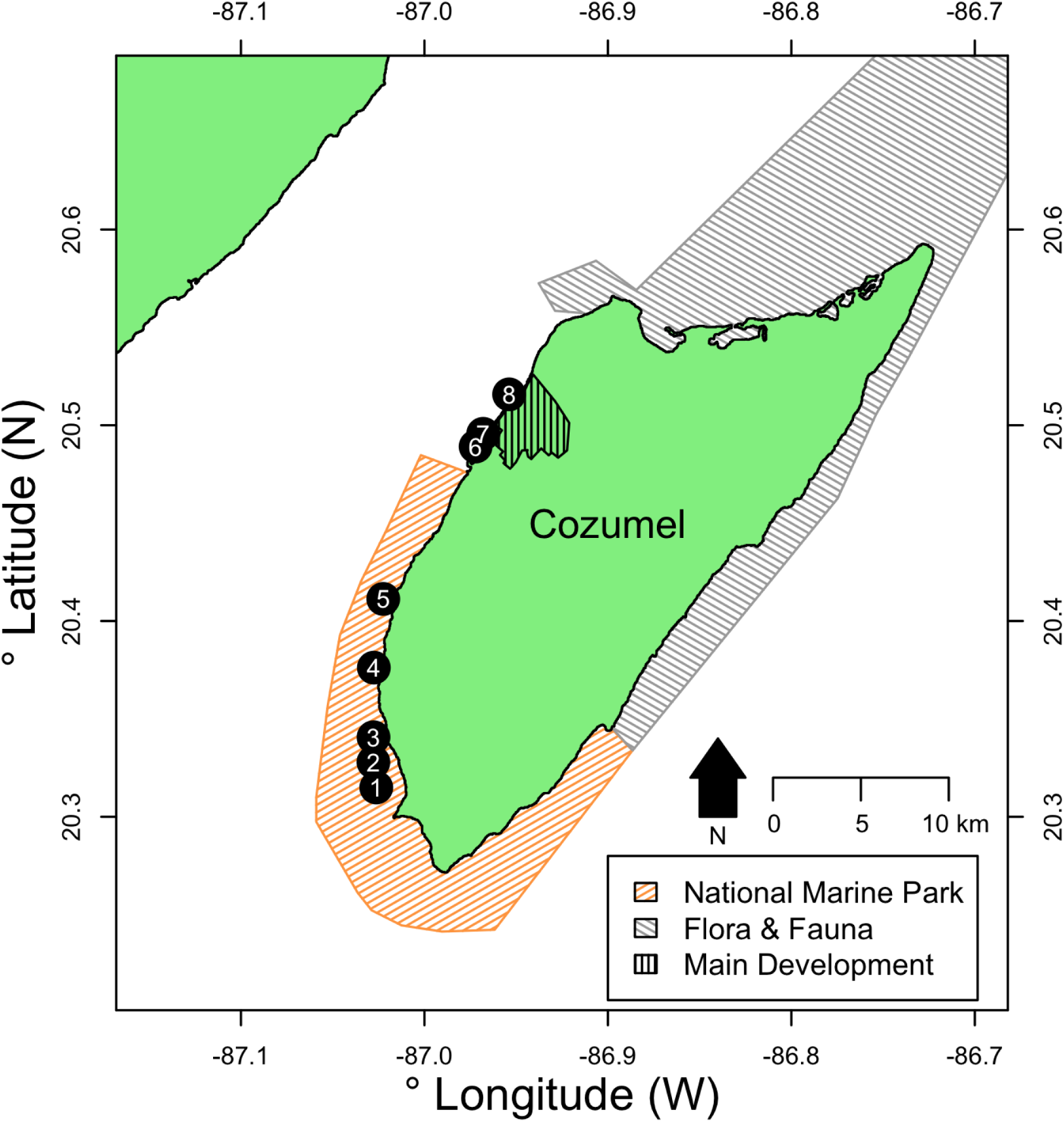
Location of survey sites relative to Cozumel and the National Marine Park and Flora & Fauna protected areas on Cozumel. Sites and their approximate distances from the main development in parenthesis were: 1 – Colombia (24.9 km), 2 – Herradura (23.3 km), 3 – Palancar Jardines (22.6 km), 4 – Santa Rosa (18.2 km), 5 – Punta Tunich (14.2 km), 6 – Villa Blanca (4.1 km), 7 – Transito Transbordador (3.3 km) and 8 – Purgatorio (0.6 km). All distances from the main development were measured from the passenger ferry terminal in the centre of town following the edge of the reef crest in Google Earth.

Surveys were conducted at eight sites around Cozumel during August 2016. Five sites were within the Cozumel National Marine Park (MPA), and three were in an area with no protection. The MPA sites were Santa Rosa, Colombia, Punta Tunich, Palancar Jardines and Herradura, and non-MPA sites were Transito Transbordador, Purgatorio and Villa Blanca outside of the MPA (Figure 1). Full GPS locations for sites are given in Electronic Supplementary Material (ESM) 1.

### Reef surveys

All surveys were conducted using open-circuit SCUBA equipment between the hours of 07:00am – 11:00am. Fish surveys were conducted using a diver-operated stereo-video system (stereo-DOV), consisting of two cameras separated by 0.8 m and with approximately 3 ^°^convergence angle filming forward along the reef (see [50] for system overview). The stereo-DOV system records two synchronised images of reef fish, allowing accurate measurements of fish length. The stereo-DOV used two GoPro Hero 4 Black cameras and a spool system with biodegradable line for measuring out each transect. Transects were 30 m in length and each separated by a 10 m interval, with four transects conducted at both 15 m (shallow) and 55 m (MCE) at each site. At the beginning of the dive the stereo-DOV operator started the cameras recording and synchronised them using a torch which was turned on and off repeatedly by the dive buddy. The cameras were then pointed downwards whilst the buddy attached the end of the biodegradable line to the reef. The stereo-DOV operator swam with the cameras down, reeling out the line, until the first marker was reached after 10 m of line. At this point the cameras were pointed forwards along the reef to record the transect. After reaching the marker indicating a further 30 m of line had been unreeled the cameras were pointed back down for 10 m before starting the next transect. This was repeated over 4 transects, with all transect start and end points, and transect intervals pre-marked on the biodegradable line.

Benthic surveys were conducted along the same survey lines following the SVS, using a GoPro Hero 4 Black camera. A planar photo quadrat was taken at the start and then at every 2.5 m intervals along the transect giving 13 quadrats per transect. When taking quadrats, the camera was held perpendicular to the reef at approximately 0.4 m above the benthos.

### Video analysis

The stereo-DOV footage was analysed using EventMeasure (v4.42, SeaGIS, Melbourne, Australia). Transects were synchronised, and all fish 2.5 m either side of the camera (5 m transect width; constrained using EventMeasure) were identified to species, or the lowest taxonomic level possible and measured from snout to the tip of caudal peduncle. From the length and species identification the biomass was estimated based on length-weight ratios from Fishbase [51], based on the equation: W=aL^b^ Where W is the weight, L is the length and a and b are given parameters for a specific species.

Photos were analysed using Coral Point Count with Excel extensions [52] to determine the percent cover of different benthic categories. Ten random points were placed on each quadrat image in CPCe, and the substrate category at each point was identified. The total number of points of each substrate category per transect was then used to calculate benthic percentage coverage for each transect. Categories were: Black Coral (Antipatharia), Hard Coral (Scleractinia), Calcareous Macroalgae, Fleshy Macroalgae, Turf Algae, Crustose Coralline Algae, Sponge, Gorgonian, Hydrozoan, Cyanobateria, and Non-Living substrate.

### Data analysis

To evaluate differences in percentage coverage of key benthic groups a Euclidian permutational analysis of variance (ANOVA) was used on mean percentage cover of each benthic group at each depth and site. To test for broader differences in benthic community assemblage based on depth, protection, and interactions between these factors, permutational multivariate analysis of variance (PERMANOVA) was used on Bray-Curtis dissimilarities of percentage cover of all benthic categories. To further explore differences in benthic community structure based on protection and depth a redundancy analysis was conducted using the function ‘rda’ in vegan [53]. This redundancy analysis was based on removing non-living substrate and standardising the percentage community composition of all living components of the community.

Commercially-important fish species were identified based on a fishbase [51] price category classification of moderate, high or very high fisheries value. Differences in fish species richness, biomass and commercially-important fish biomass were identified using ANOVA fitting depth and protection as factors. Residual plots were checked after model fitting to ensure model assumptions were not violated. Models were simplified to remove non-significant factors or interactions based on minimising the Akaike information criterion (AIC). To identify differences in commercially-important fish, we totalled the commercially-important species biomass by family and used permutational ANOVA to test for effects of depth and protection. We followed Langlois et al. [54] to use kernel density estimates to compare length distributions between fish surveyed within and outside the protected area. Bandwidths were selected using the Sheather-Jones selection procedure [55] within the ‘dpik’ function in the ‘KernSmooth’ package [56]. Differences in the length distributions were then tested using the permutational ‘sm.density.compare’ function in the R package ‘sm’ [57].

All permutational ANOVAs and PERMANOVAs were fitted using the ‘adonis’ function in vegan [53]and run for 99999 permutations. All analysis was conducted in R [58].

## Results

### Benthic communities

We identified differences in benthic communities based on both protection status and depth, with the significant interaction between protection and depth indicating that the effect of protection changes based on depth (Table 1). We found greater hard coral cover on shallow reefs inside the protected area (8.5 ± 2.9 *%* cover; mean ± SE) than outside (0.5 ± 0.1 %), and greater gorgonian coverage on MCEs inside the protected area (7.1 ± 1.6 %) than outside (1.6 ± 0.7 %) (Figure 2). No other significant differences were detected between percentage cover of major groups such as sponges, macroalgae and non-living substrate between areas of the same depth based on protection (Figure 2). There were major differences in benthic cover between shallow reefs and MCEs, with all surveyed Cozumel MCEs existing as continuous reef systems dominated by sponges and calcareous macroalgae (mostly *Halimeda)*, with black corals present and very little of the benthos covered by non-living substrates (Figure 2B). In contrast, the shallow reefs of Cozumel were characterised by areas of reef separated by patches of sand resulting in higher non-living benthic cover (Figure 2A). A full list of hard coral and black coral species identified at each depth is contained in ESM 2.

**Table 1.** Benthic PERMANOVA testing for differences in benthic community structure between different protection types, depths and sites, and the interactions between them.

| Source | DF | Mean Square | Pseudo-F | *p* |
| --- | --- | --- | --- | --- |
| Protection | 1 | 0.5 | 9.8 | <0.001 |
| Depth | 1 | 0.9 | 16.1 | <0.001 |
| Site | 6 | 0.3 | 4.8 | <0.001 |
| Protection:Depth | 1 | 0.4 | 7.7 | <0.001 |
| Depth:Site | 6 | 0.2 | 4.0 | <0.001 |
| Residuals | 48 | 0.1 |  |  |
| Total | 63 |  |  |  |

**Figure 2.**
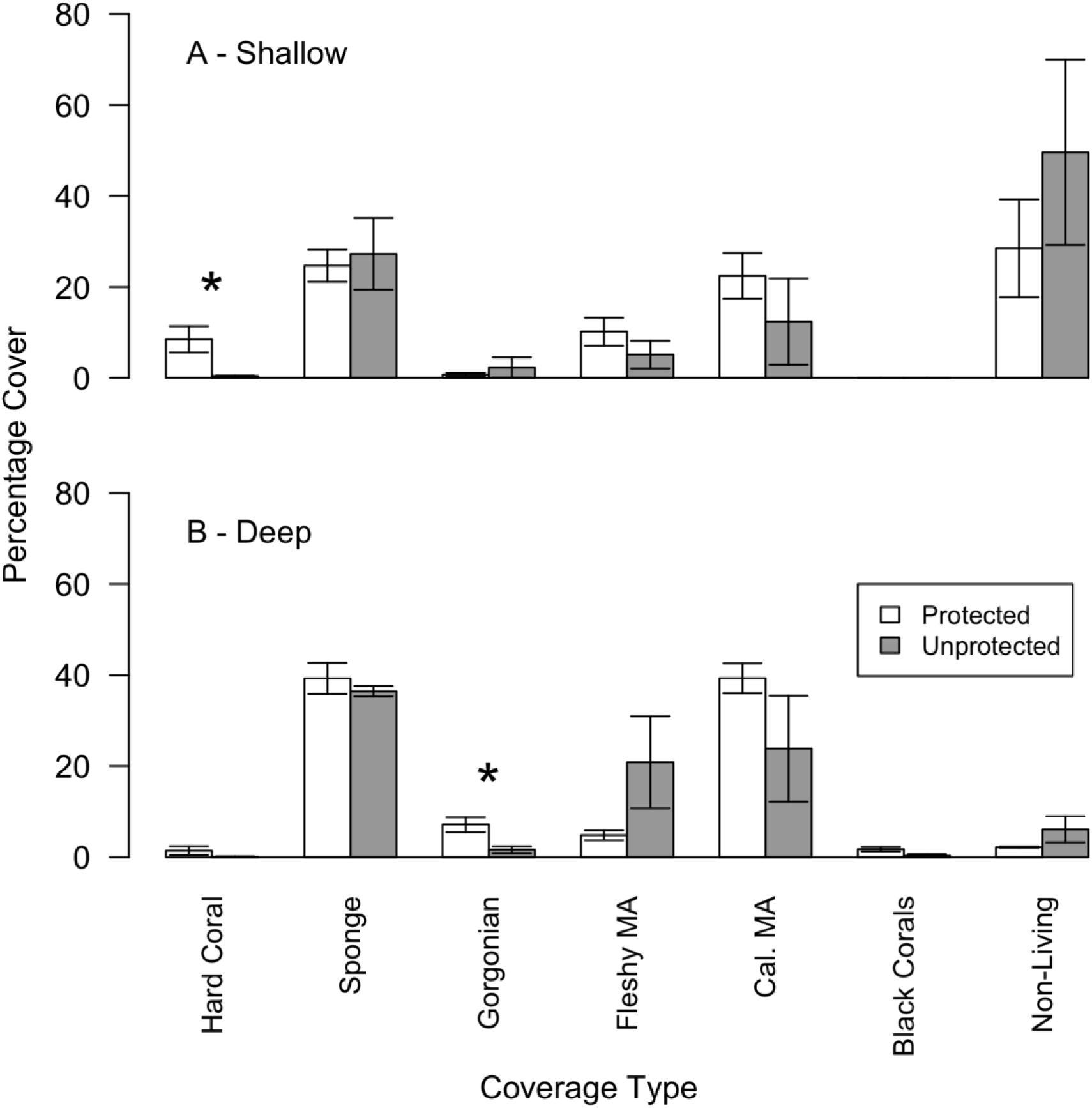
Percentage cover of broad benthic groups on (A) shallow reefs at 15 m and (B) MCEs at 55 m around Cozumel. Error bars represent one standard error. Significantly different coverage (*p*<0.05) between protected and unprotected areas was tested using a permutational ANOVA and indicated with a ‘*’.

To further explore differences in benthic ecological communities between sites within the protected area and those outside we conducted a redundancy analysis (RDA) of the benthic coverage data after removing non-living benthic groups and recalculating percentages. In the shallows we found that two of our three sites without protection were correlated with higher sponge cover, while the other site without protection had higher gorgonian and hydroid cover (Figure 3A). The highest hard coral cover was associated with two of the protected sites, Palancar Jardines and Herradura, at 15.7 ± 6.9 % and 14.4 ± 2.3 % cover respectively. While the three sites without protection had the lowest hard coral cover at 0.6 ± 0.6 % (Purgatorio), 0.2 ± 0.2 % (Transito Transbordador) and 0.6 ± 0.4 % (Villa Blanca). On MCEs, protected sites were associated with greater gorgonian, black coral and crustose coralline algae cover (Figure 3B). Interestingly, some sites which clustered close together in the RDA analysis in the shallows also did so on MCEs, for example, outside the protected area Transito Transbordador and Purgatorio, and inside the protected area Palancar Jardines and Herradura. This suggests similar environmental or anthropogenic processes may be driving benthic communities on shallow reefs and MCEs. In addition to being associated with higher hard coral cover in the shallows, both Palancar Jardines and Herradura were associated with higher hard coral cover on MCEs (Figure 3B), with Herradura having the highest hard coral coverage we observed on Cozumel MCEs at 5.1 ± 2.0 %. Black corals were recorded at all five MCEs within the protected area, but only at the Purgatorio MCE outside the marine park. However, overall recorded black coral coverage was low, with 3.0 ± 1.2 % at Palancar Jardines and 2.9 ± 2.9 % at Santa Rosa, the two sites with the greatest coverage.

**Figure 3.**
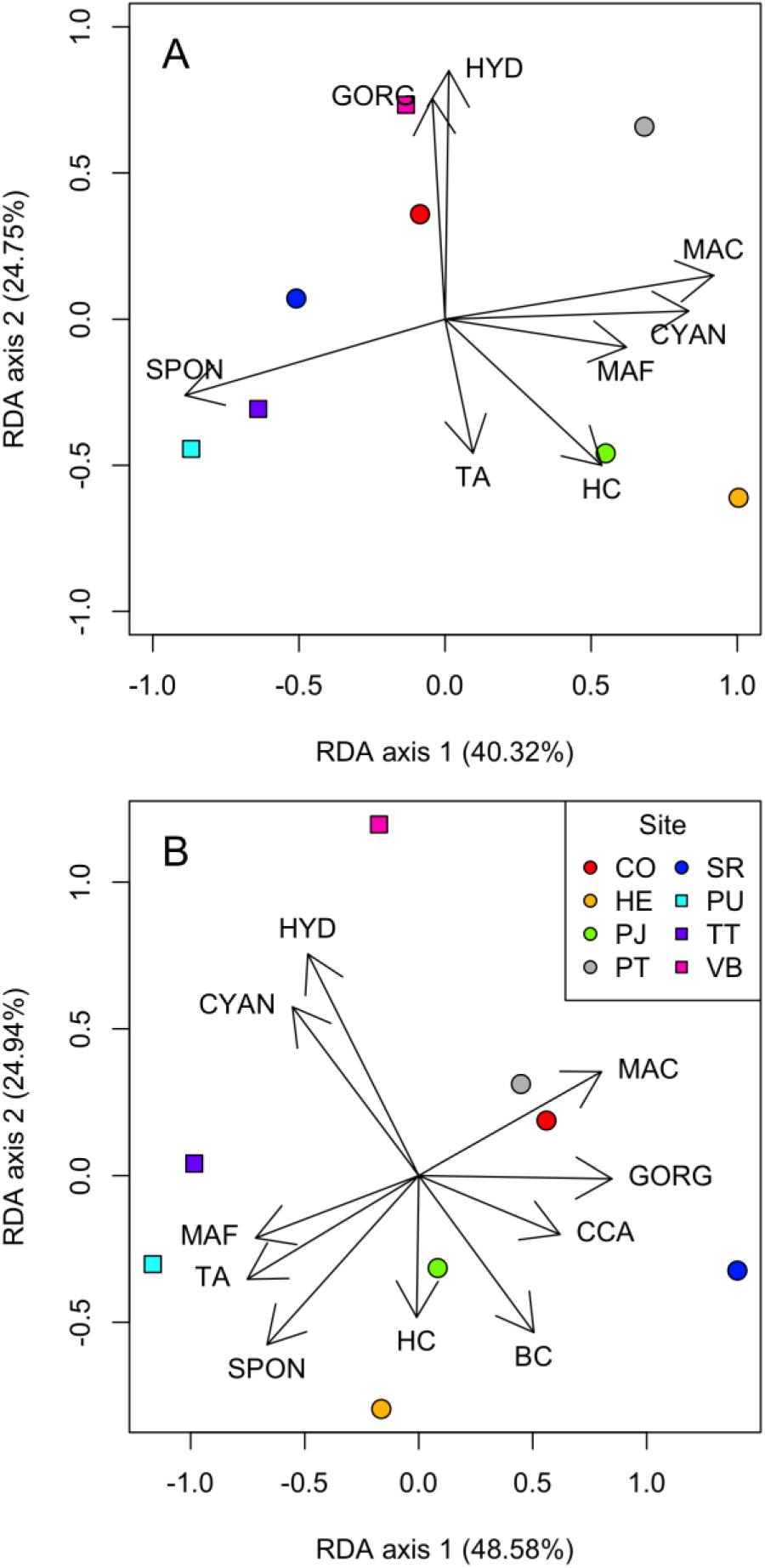
Redundancy analysis of the benthic coverage data standardised to remove non-living benthic cover for (A) shallow reefs at 15 m, and (B) MCEs at 55 m. Variation explained by each axis is indicated in parenthesis on the axis labels. The length and direction of the arrows corresponds to increasing cover of benthic categories at sites located in that region of the plot. Benthic categories were: BC – black coral, CCA – crustose coralline algae, CYAN – cyanobacteria, GORG – gorgonian, HC – hard coral, HYD – hydrozoan, MAC – calcareous macroalgae, MAF – fleshy macroalgae, SPON – sponge, and TA – turf algae.

### Fish communities

No difference in fish species richness was identified between shallow reefs located inside and outside the protected area or between MCEs located inside and outside the protected area (Figure 4A). However, fish species richness was greater on shallow reefs than MCEs (F_1,13_=22.8, *p*<0.001), with a mean shallow reef fish species richness of 12.4 ± 0.7 species per 150 m^2^ in contrast to 7.6 ± 0.6 mean species richness per 150 m^2^ on MCEs. Overall, we recorded 80 fish species on Cozumel reefs in this study, with 39 species (48.8 %) only recorded on shallow reefs, 7 species (8.9 %) only recorded on MCEs and 34 species (42.5 %) recorded on both shallow reefs and MCEs. The full list of which species were recorded at one or both depths is available in ESM 3.

**Figure 4.**
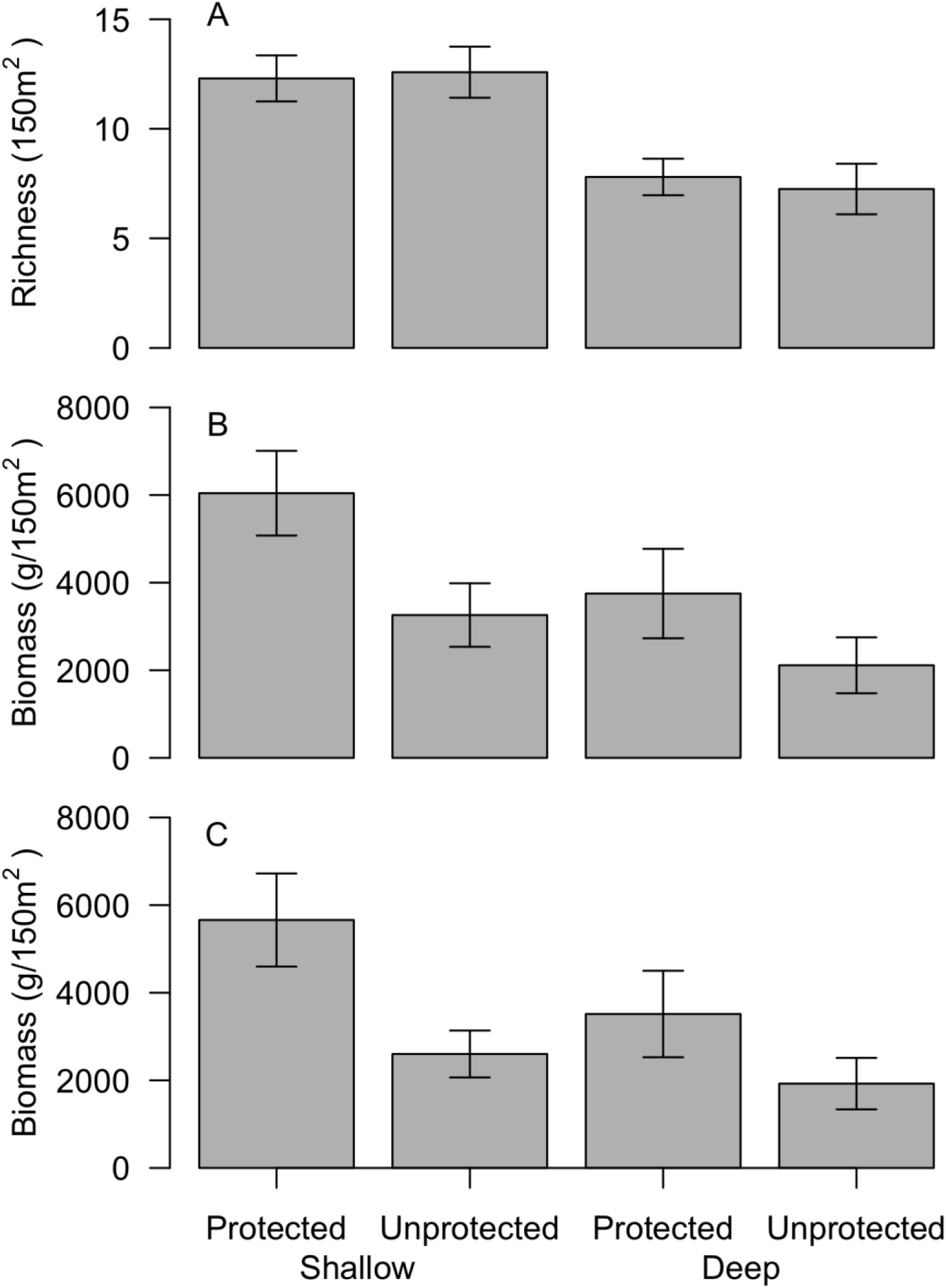
Comparisons of reef fish communities for shallow (15 m) and mesophotic (55 m) for (A) species richness, (B) all fish biomass, and (C) commercially-important fish biomass. Error bars indicate one standard error.

We detected weak effects of protection status on both overall fish biomass (F1, _13_=5.1, *p*=0.04) and commercially-important fish biomass (F_1,13_=5.5, *p*=0.04), with greater fish biomass associated with sites within the protected area on both shallow reefs and MCEs (Figure 4B, 4C). We found no significant interaction between depth and protection (so removed this interaction from the model during simplification) or effect of depth (shallow vs MCE) on overall fish biomass (F_1,13_=3.9, *p*=0.07; Figure 4B) or commercially-important fish biomass (F_1,13_=2.8, *p*=0.12; Figure 4C). However, during model simplification for both overall fish biomass and commercially-important fish biomass we found that removing depth from the model resulted in a greater model AIC value than retaining it (Model AIC for overall fish biomass: 293.66 without depth versus 291.47 with depth included; commercially-important fish biomass: 293.66 without depth versus 291.98 with depth included), suggesting that differences with depth may affect reef fish biomass.

To identify which fish families might be driving these patterns, and to investigate the potential depth refuges for important fisheries species, we grouped all commercially-important fish species by family and compared their biomass inside and outside the marine park, and on shallow reefs and MCEs using a permutational ANOVA (Table 2). We found no commercially-important fish families showed interactions between depth and protection, or protection effects (Table 2). Commercially-important species, comprising four fish families, biomass was affected by depth, however the effect of depth was not consistent between families. Three families showed reduced biomass on MCEs compared to the shallows, these were (percentage decline in biomass for shallow reefs vs. MCEs in parenthesis): Acanthuridae (74.9 %), Haemulidae (96.0 %) and Mullidae (100.0 %). While Pomacanthidae showed a 396.1 % increase on MCEs compared to shallow reefs.

**Table 2.** Biomass of commercially-important fish species grouped by family from inside and outside the marine park on shallow reefs and MCEs. Depth and protection effects were tested using a permutational ANOVA, with significant effects (*p*<0.05) highlighted in bold.

| Family | Shallow reefs |  |  |  | MCEs |  |  |  | Depth effect |  | Protection effect |  | Depth:Protection interaction |  |
| --- | --- | --- | --- | --- | --- | --- | --- | --- | --- | --- | --- | --- | --- | --- |
|  | Inside Park |  | Outside Park |  | Inside Park |  | Outside Park |  |  |  |  |  |  |  |
|  | Mean biomass (g/150 m<sup>2</sup>) | SE | Mean biomass (g/150 m<sup>2</sup>) | SE | Mean biomass (g/150 m<sup>2</sup>) | SE | Mean biomass (g/150 m<sup>2</sup>) | SE | Pseudo-F | *p* | Pseudo-F | *p* | Pseudo-F | *p* |
| Acanthuridae | 845 | 249 | 1163 | 330 | 239 | 91 | 247 | 122 | 11.65 | **<0.01** | 0.56 | 0.47 | 0.51 | 0.49 |
| Balistidae | 336 | 177 | 170 | 65 | 258 | 225 | 181 | 141 | 0.05 | 0.78 | 0.37 | 0.56 | 0.05 | 0.81 |
| Carangidae | 1083 | 913 | 0 | 0 | 61 | 41 | 64 | 39 | 1.09 | 0.38 | 0.79 | 0.44 | 0.79 | 0.45 |
| Haemulidae | 468 | 333 | 429 | 215 | 0 | 0 | 48 | 25 | 3.64 | **0.02** | 0.00 | 0.99 | 0.03 | 0.90 |
| Kyphosidae | 51 | 51 | 0 | 0 | 0 | 0 | 0 | 0 | 0.94 | 1.00 | 0.56 | 1.00 | 0.56 | 1.00 |
| Labridae | 163 | 120 | 0 | 0 | 6 | 6 | 0 | 0 | 1.61 | 0.33 | 1.13 | 0.26 | 0.96 | 0.48 |
| Lutjanidae | 817 | 345 | 72 | 72 | 537 | 534 | 0 | 0 | 0.24 | 0.56 | 2.28 | 0.15 | 0.06 | 0.78 |
| Malacanthidae | 10 | 10 | 0 | 0 | 0 | 0 | 0 | 0 | 0.94 | 1.00 | 0.56 | 0.37 | 0.56 | 1.00 |
| Monacanthidae | 30 | 30 | 20 | 12 | 0 | 0 | 0 | 0 | 1.73 | 0.21 | 0.07 | 0.91 | 0.07 | 0.98 |
| Mullidae | 0 | 0 | 18 | 1<br>2 | 0 | 0 | 0 | 0 | 2.38 | <0.<br>01 | 3.96 | 0.<br>0<br>5 | 3.96 | 0.0<br>5 |
| Ostracii<br>dae | 56 | 5<br>6 | 0 | 0 | 0 | 0 | 0 | 0 | 0.94 | 1.0<br>0 | 0.56 | 0.<br>3<br>8 | 0.56 | 1.0<br>0 |
| Pomaca<br>nthidae | 228 | 1<br>4<br>7 | 48 | 2<br>4 | 614 | 22<br>0 | 109<br>8 | 5<br>5<br>6 | 5.94 | 0.0<br>3 | 0.32 | 0.<br>5<br>8 | 1.52 | 0.2<br>4 |
| Scaridae | 920 | 3<br>0<br>7 | 661 | 3<br>9<br>9 | 326 | 14<br>8 | 224 | 8<br>2 | 4.14 | 0.0<br>6 | 0.44 | 0.<br>5<br>3 | 0.08 | 0.7<br>8 |
| Scorpae<br>nidae | 0 | 0 | 0 | 0 | 42 | 42 | 46 | 4<br>6 | 1.90 | 0.3<br>4 | 0.01 | 0.<br>7<br>5 | 0.01 | 0.8<br>7 |
| Serranid<br>ae | 63 | 6<br>1 | 21 | 1<br>1 | 143<br>2 | 13<br>88 | 16 | 1<br>0 | 0.91 | 0.7<br>1 | 0.62 | 0.<br>5<br>7 | 0.55 | 0.7<br>0 |
| Sphyrae<br>nidae | 590 | 5<br>9<br>0 | 0 | 0 | 0 | 0 | 0 | 0 | 0.94 | 1.0<br>0 | 0.56 | 0.<br>3<br>7 | 0.56 | 1.0<br>0 |

We tested fish length distributions, comparing inside and outside the protected area, finding that in shallow reefs outside the protected area a greater proportion of the fish are of small (>200 mm) body length (Figure 5A). This pattern is even more extreme when considering only commercially-important species on unprotected shallow reefs, with a large peak in fish body lengths between 100-250 mm, and few individuals bigger than 300 mm (Figure 5C). While protected shallow reefs share having many fish in the 100-250 mm range, there are more fish with greater body lengths in the 250-400 mm range (Figure 5C). In contrast, on MCEs there are less clear differences between fish length distributions inside and outside the protected area. While there are statistically significant differences in the length distribution for all recorded MCE fish, this appears to be driven by differences in the proportion of smaller fish in the 0-100 mm length range with larger bodied fish showing similar proportions (Figure 5B). When specifically comparing commercially-important fish on MCEs, we found no difference in the fish length distributions based on protection status (Figure 5D). In general, we recorded few large fish on reefs at both depths and protection types around Cozumel, with only 10 individuals >500 mm length out of the 2,599 recorded fish. These were individuals of: *Caranx latus, Mycteroperca bonaci, Ocyurus chrysurus, Pomacanthus arcuatus* and *Sphyraena barracuda*.

**Figure 5.**
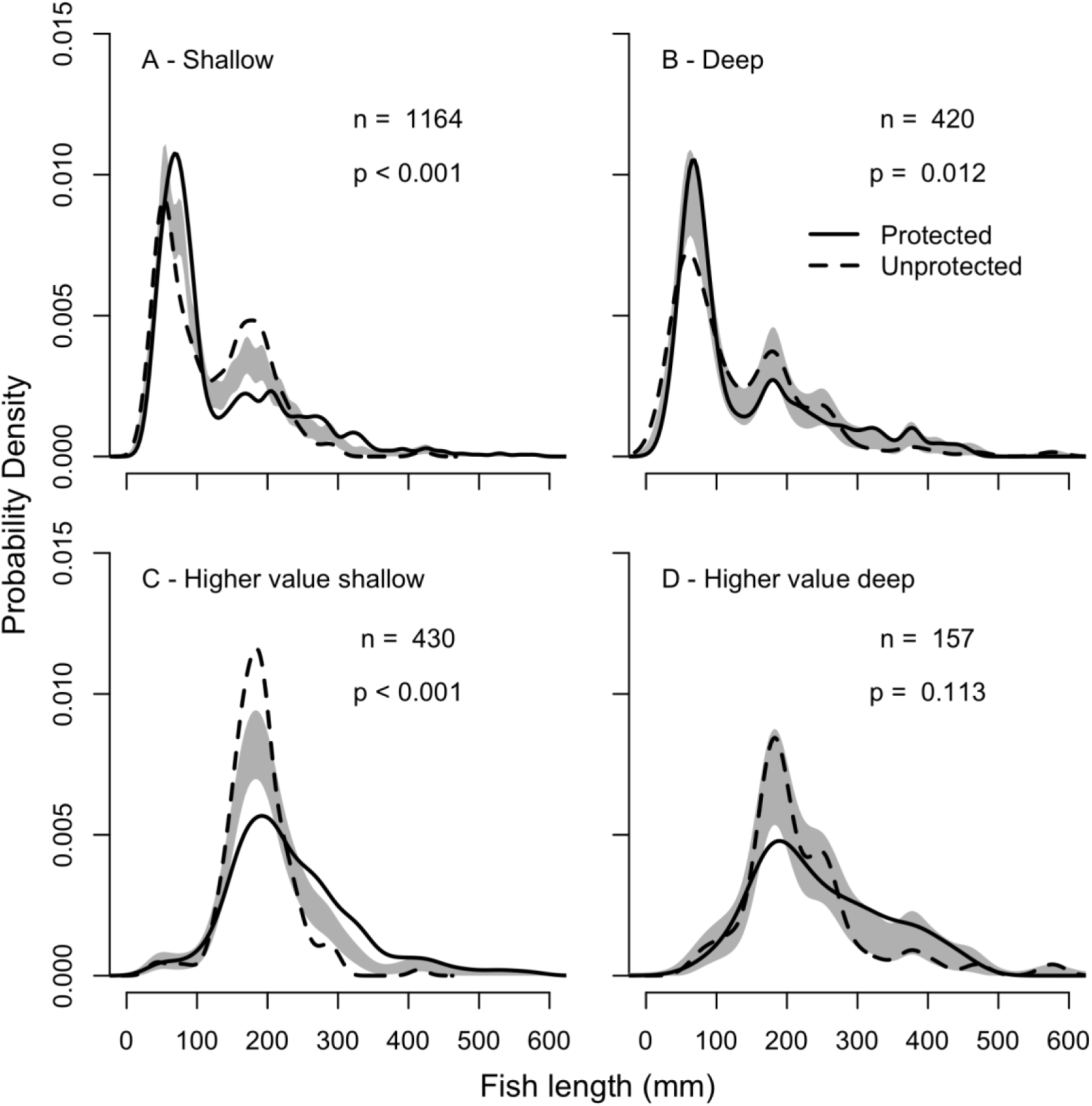
Fish length distributions for all fish species for (A) shallow reefs, (B) MCEs, and for commercially-important fish species only for (C) shallow reefs and (D) MCEs. The grey shaded area indicates one standard error either side of the null model of no difference in length distribution based on protection. n=number of fish.

## Discussion

In order to test whether MCEs act as deep reef refuges, two aspects need to be considered: (i) the extent MCEs are protected from disturbances affecting shallow reefs, and (ii) evidence that MCEs could help repopulate shallow areas following disturbance [20]. Our results show that Cozumel MCEs benthic communities appear similar between sites within the protected area and areas adjacent to large shallow reef impacts. This supports the idea that MCEs have the potential to serve as refuge for benthic species. However, we identified that most hard coral species found on shallow reefs decrease in abundance or are absent on MCEs, suggesting that MCEs may have limited ability to aid shallow reef hard coral recovery. In contrast, we found 42.5 % of fish species recorded on both shallow reefs and MCEs, including many commercially-important fish species. Our results therefore indicate that MCEs may play a role in supporting fish populations. However, regardless of protection we found few large-body fishes (>500 mm), which were nearly absent at all studied sites.

### Differences between inside and outside MPA for shallow reefs and MCEs

We tested whether reefs within the MPA were similar to those outside. We found that while the MPA had higher hard coral cover for shallow reefs, the main difference between MCEs inside and outside the protected area is the higher abundance of gorgonians inside. Hard corals represent a major component of the benthic community providing structural habitat in the shallow areas. Previous research has reported large declines in shallow reef hard coral cover in the area without protection on Cozumel, including at one of our study sites Villa Blanca [38]. At Villa Blanca hard coral cover declined from 44 % in 1995 to 4 % in 2005 [38], which is more severe than declines recorded within the protected area during this time [59]. We recorded current hard coral cover at Villa Blanca at <1 % suggesting that further declines have occurred. This unprotected area is adjacent to Cozumel town with multiple cruise ships, passenger and car ferries passing over and docking adjunct to the reef daily. In addition, development of a large cruise ship terminal appears to have severely affect shallow reefs [38,39].

In general, reefs outside the protected area were dominated by non-living components (e.g. discarded artificial structures and sand). In contrast, we found much greater hard coral cover on shallow reefs inside the protected area (8.5 ± 2.9 % cover; mean ± SE), this is similar to estimates from recent Cozumel reef monitoring surveys inside the protected area [59,60]. Even within the protected area however, shallow reef communities exist as a series of built up reefs separated by patches of sand, and so have a large proportion of non-living benthic cover. The percentage of non-living benthic cover was not different on shallow reefs between the MPA and areas outside, we think this maybe partly because of the areas surveyed. With more replicates/larger surveyed area it is possible that more patterns would have been detectable, and we recommend this for future studies.

Regardless of protection and location, all observed Cozumel MCEs were continuous reefs with the main structural habitat complexity provided by calcareous macroalgae, sponges, gorgonians, and black corals. While hard corals were present on MCEs, these were at low abundance. There was no difference between sites inside and outside the MPA on any benthic community component surveyed except gorgonians. Gorgonian abundance was greater in the protected area (7.1 ± 1.6 %) than unprotected sites (1.6 ± 0.7 %). It is not clear what drives these patterns, as it has previously been suggested that gorgonians are more resilient to disturbance impacts and other environmental factors than many other reef organisms such as hard corals [61,62]. However, the lack of hard corals on MCEs combined with high densities of gorgonians may mean that gorgonians are a better indicator of MCE state [12]. In this context our results would suggest that the disturbance associated with Cozumel town and the associated boats is likely to be affecting benthic communities on MCEs.

Biomass, on both shallow reefs and MCEs, was higher within the protected area than outside for all fish species, and also for commercially-important fish species. Despite the higher fish biomass within the protected area than outside, Cozumel shallow reef fish biomass within the protected area is considered low for the region [60]. This suggests that shallow sites outside the protected area are even more severely depleted. These shallow reef findings are further supported by the fish length distributions, showing fewer large fish on shallow reefs outside the protected area, particularly those of higher commercially-important. This contrasts with fish length distribution comparisons for MCEs, where there was no difference for commercially-important fish between sites within and outside the MPA. While this potentially suggests a depth refuge for larger fish on MCEs outside the protected area, this finding must be treated with caution. Fewer commercially-important fish were measured on MCEs than shallow reefs (157 versus 430), reducing power to discern differences based on protection on MCEs. In addition, the length distributions for commercially-important fish on MCEs comparing protection status looks very similar in shape to those shown for comparisons based on protection status on shallow reef commercially-important fish (Figure 5C-D). This suggests that further work is required to establish whether there are differences in length distributions based on protection on MCEs.

Regardless of protection and depth we found only 10 individual fish >500 mm length out of the 2,599 recorded fish. This suggests a general absence of large predatory fish from the reefs of Cozumel, and is consistent with other studies on shallow reefs and MCEs facing fisheries pressure within the Mesoamerican Barrier Reef region. For example, surveys conducted on almost 150 Mesoamerican Barrier Reef shallow sites found that large groupers (>400 mm) were highly scarce, present in only 11% of locations [60]. While studies on MCEs on the southern Mesoamerican Barrier Reef have revealed increased fish body size on MCEs compared to shallow reefs, suggesting possible refuges, there were still limited numbers of larger predatory fish found [18]. However, other studies have identified that Caribbean MCEs do appear to be acting as refuges for historically overfished large predatory species such as sharks and groupers [19,63].

Care must be taken when interpreting comparisons between our protected sites and our unprotected area. Unfortunately, because of the location of the National Marine Park on the south west coast and the unprotected area adjacent to Cozumel town on the west coast, it has not been possible to clearly disentangle effects of protection from a geographical gradient along the Cozumel coast. Previous research has repeatedly shown more severe declines in shallow reef condition in the area without protection than has been recorded for the protected area [38,39,60]. This decline in shallow reef health outside the protected area has been attributed to the close proximity of shoreline development and the large population impact because of Cozumel town [38,60] combined with large port developments adjacent to the reef [39]. Our sites therefore exist on a gradient of increasing distance from the largest human settlement. Other processes can also be identified along this geographical gradient. For example, currents predominantly flow from south to north along the west coast of Cozumel [64]. Currents can influence water quality and correlate with both benthic and fish community structure [65,66]. However the greatest effects of currents on reef communities have been recorded in lagoons where water flow is restricted [66,67]. This suggests that while the current flowing past the reefs of Cozumel are likely to affect communities, this current gradient is unlikely to be the primary drivers of decline in for reefs in the more northern unprotected area.

### Community ecology across shallow reefs to MCEs around Cozumel

All surveyed MCEs were located on steep slopes as extensions of the shallow reef community. This characteristic reduces the light levels available to benthic organisms rapidly with increased depth [7,12]. MCEs had lower hard coral cover than the shallows, which is consistent with previous preliminary observations of MCEs around Cozumel [61,68]. For example, Dahlgren [61] reports that hard coral dominated reefs ended at approximately 30 m in at the sites within the protected area, including two of our study sites: Colombia and Santa Rosa. While Günther [68] conducted surveys to 40 m depth and reports that the deeper slopes in the 40-50 m range of Cozumel are dominated algae with large sponges and octocorals present. They also report small isolated hard coral colonies present of mostly *H. cucullata, P. astreoides* and *E. fastigiata*. Interestingly, while quantitative data broken down by site and depth is not available from these earlier studies, our results appear to suggest that unlike shallow reefs, MCEs on Cozumel have not changed much in broad benthic composition. For example, we observed high presence of macroalgae, sponges and octocorals, as well as small colonies of *H. cucullata* present. This supports the idea that MCEs by virtue of their depth have provided some protection, and the main benthic communities that provide habitat and supports many other organisms are macroalgae, sponges, gorgonians and black corals.

Surprisingly we did not find a strong effect of depth on fish biomass. However, our model simplification based on AIC suggested that depth did have useful explanatory power when considering fish biomass. Decreasing fish biomass with increasing depth has been documented on the southern Mesoamerican Barrier Reef [18], and also at other locations in the Caribbean such as Curaçao [69] and Puerto Rico [19]. It is not clear why we did not observe this pattern, though while it is possible this could be caused by fisheries pressure on shallow reefs removing shallow reef fish biomass. While Figure 4 does not show a significant difference in biomass based on depth, it is suggestive that with greater statistical power a difference may be detectable. Recent work conducted on the Mesoamerican Barrier Reef has also suggested that stereo-DOV surveys may bias against smaller fish on MCEs compared to other fish survey techniques [24], though it is not clear whether this is through diver avoidance or reduced ability to discern fish on videos with lower levels of lighting. However, even if some smaller fish were missed on transects, these individuals will likely have lower contribution to overall fish biomass and so are unlikely to drive patterns in overall fish biomass with depth. We recommend more transects, of larger areas should be conducted in future studies to examine fish biomass patterns with depth in Cozumel reefs.

While we detected no overall difference in fish biomass between shallow reefs and MCEs, for several commercially-important fish species patterns were apparent. Biomass of commercially-important Acanthuridae, Haemulidae and Mullidae declined with increased depth. Patterns of decline in herbivorous fish biomass has been widely observed on MCEs in the western Atlantic [18,19,70], so declines in herbivorous Acanthuridae are not surprising. However, previous studies have identified species of Haemulidae as indicators of Caribbean MCEs [19], and Haemulidae have been observed on Mersoamercian Barrier Reef MCEs in Belize [71]. Additionally, despite only recording Mullidae on shallow reefs in our surveys, in Belize they have been observed >100 m on MCEs [71]. In contrast, commercially-important Pomacanthidae increased in biomass on MCEs, likely caused by the increased cover of sponges as many Pomacanthidae species are spongivores [72].

The sites furthest south (Palancar Jardines and Herradura in our study) had higher shallow hard coral cover than the other sites inside the protected area further north and unprotected sites. These furthest south sites also had the highest hard coral cover on MCEs, suggesting that factors driving these hard coral cover in the shallows may also be influencing MCEs. Both of these sites are furthest away from the main area of development on Cozumel, and the first reefs that currents pass over along the coast of Cozumel. The influence of both distance from settlement and current strength should be investigated in future studies.

### Integrating MCEs into current MPA management

Our results highlight that MCEs contain highly developed benthic communities with many fish species previously reported on shallow reefs associated with them. While there is some evidence that they may be buffered from some of the disturbances affecting unprotected shallow reefs; our results also indicate that they contain unique benthic assemblages that can benefit from protection. When designing and implementing reef management plans, the whole reef ecosystem should be considered including MCEs [17]. Previous examples suggest that in places where coral reef management is already in place for shallow areas, incorporation of MCEs does not need to be complex [9,17].

Recent work has highlighted the refuge role that MCEs can play for invasive lionfish in the Caribbean [73], which in areas with shallow reef culling can still leave large lionfish abundances on MCEs [74]. On Cozumel there is widespread shallow lionfish culling by the recreational dive community and fishers, and as would be expected with sustained culling pressure we did not observe any lionfish on our shallow fish transects. We only observed two individual lionfish on our MCE transects, one at Villa Blanca and one at Herradura. Therefore, despite large lionfish refuges from culling being reported on MCEs in the southern Mesoamerican Barrier Reef [74], MCEs on the west coast of Cozumel do not appear to have a similar lionfish refuge role.

Overexploitation of shallow reef fisheries combined with new technology has been suggested to lead to expansion of fisheries to MCEs [21,24]. While in some areas of the Caribbean, MCEs have been highlighted as refuges for commercially-important fish species [19,69] our results do not support this view for Cozumel. In Cozumel, the low abundance and biomass of commercially-important fish has been well documented for the shallow areas since 2008 [59,60]. Yet current annual monitoring assessments in Cozumel are only conducted to a maximum depth of 15 m [59] leaving a large knowledge gap on deeper reefs. The current Cozumel management plan states that the National Marine Park extends to the 100 m isobath [43]. Despite this, there is no explicit acknowledgment of MCEs in the management plan and the plan implies that reef habitat does not extend beyond 30 m depth [43]. Therefore, this study emphasises the need to better incorporate deeper reefs into protected area, including implementation of fisheries and harvesting regulations.

## Conclusion

This study provides a first quantitative characterisation of MCEs around Cozumel, and compares them with adjacent shallow reefs and within and outside a protected area. We identified differences in benthic communities and fish communities between sites inside and outside the protected area, suggesting that MCEs can be affected by adjacent coastal development. Our study highlights the need to integrate MCEs in current reef management plans since they are a continuation of shallow coral reefs containing both unique species as well as many threatened and commercially-important shallow reef species.

## Acknowledgments

We wish to thank Kiragu Mwangi and Stuart Paterson (Conservation Leadership Programme) for providing support and guidance through the development of this project. Additional thanks to Alex David Rogers (University of Oxford) for providing technical advice for the project, and to Cristopher Gonzalez and Blanca Quiroga (CONANP) for assisting with literature from Cozumel reefs management.

## Ethics

This study conducted video observations of fish and benthic communities on reefs around Cozumel, Mexico. No ethical permission was required to undertake this work.

## Permission to carry out fieldwork

Permission to carry out fieldwork was granted to EG by the Comisión Nacional de Áreas Naturales Protegidas (CONANP) Dirección del Parque Nacional “Arrecifes de Cozumel”. Permit number: F00.9/DPNAC/305–16 F00.9.DRPYCM.00778/2016

## Data Availability

All raw data and R code for analysis will be made available prior to peer review.

## Competing Interests

We have no competing interests.

## Authors’ Contributions

EG and DAAB designed the study. EG, MJAG, GW and DAAB conducted the fieldwork and video/photo analysis. DAAB conducted the statistical analysis. EG, MJAG, GW and DAAB wrote the manuscript and critically revised it. All authors gave final approval for publication.

## Funding Statement

Funding for this study was provided by Arcadia through the Conservation Leadership Programme (coordinated by Flora and Fauna International, BirdLife International and Wildlife Conservation Society). DAAB is funded by a Fisheries Society of the British Isles PhD studentship. The funders did not have any role in the study design, data collection and analysis, decision to publish, or preparation of the manuscript.

---

**ESM 1.**
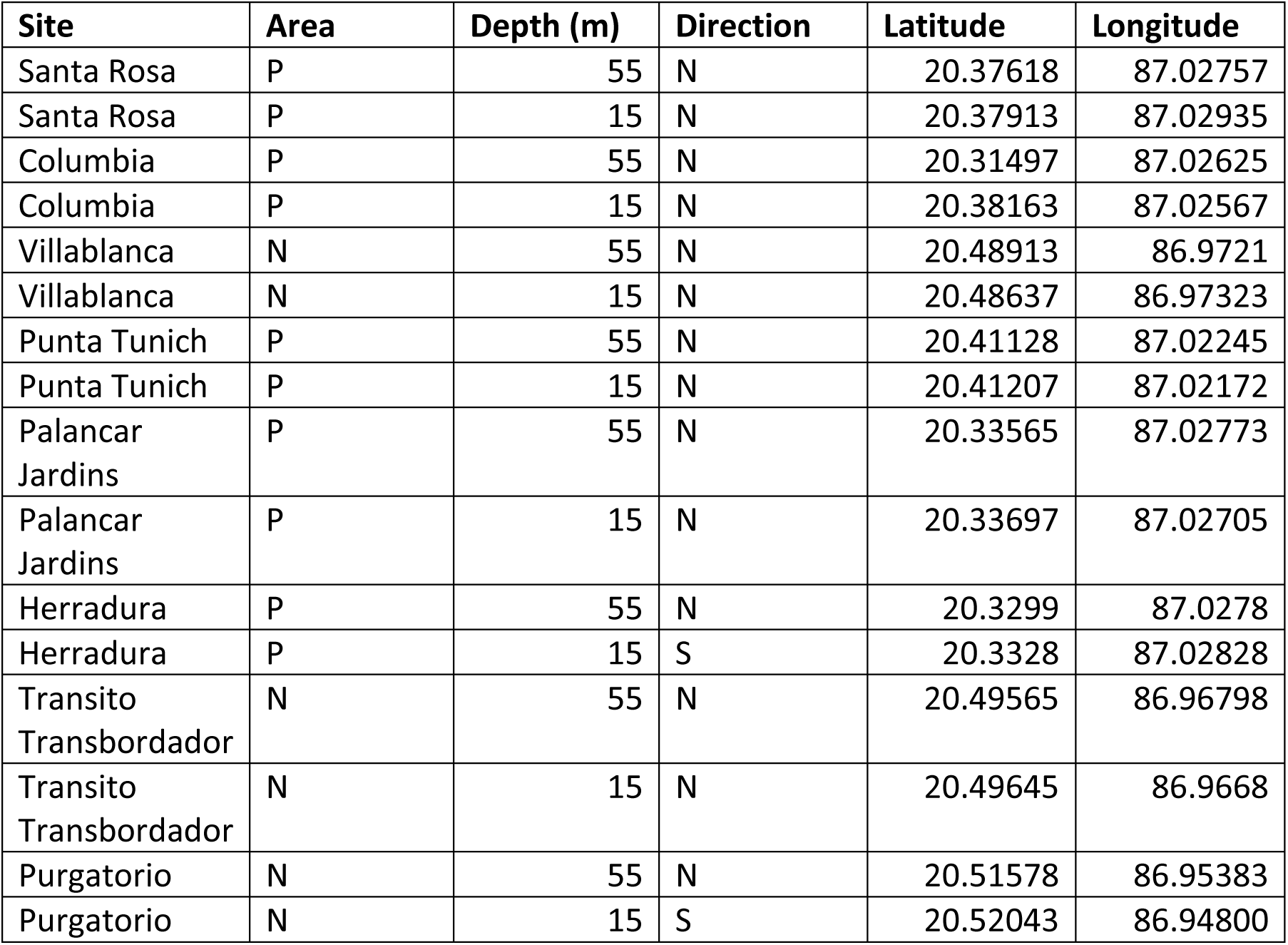
Study site GPS locations. Area indicates whether within the National Marine Park (P) or in an unprotected area (N). Direction indicates whether transects were conducted following the reef depth contour broadly north (N) or south (S) from the GPS location. All GPS points given in WGS84 format.

**ESM 2.**
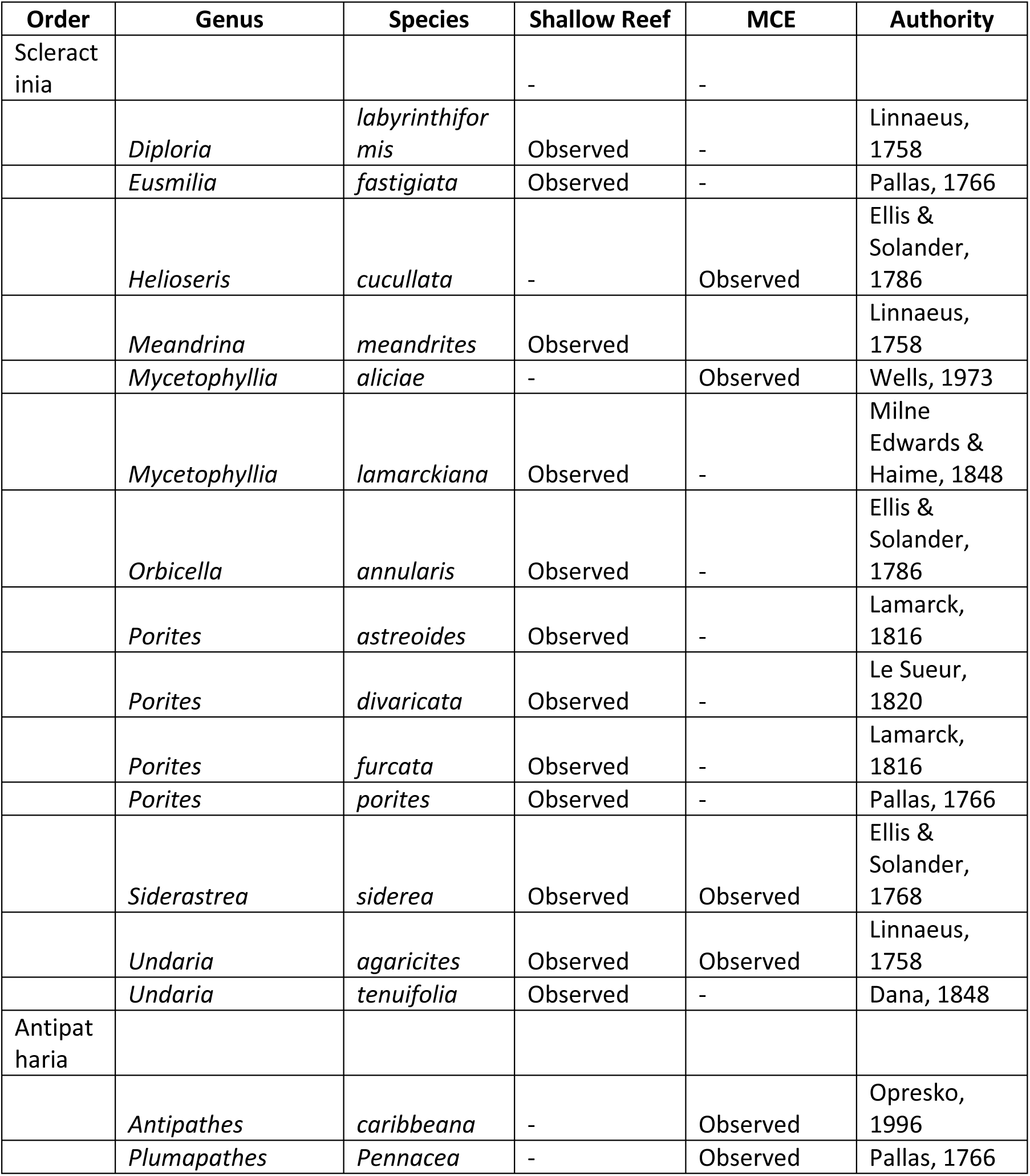
Hard coral species (Scleractinia) and black coral species (Antipatharia) observed on shallow reefs (15 m) and MCEs (55 m) at surveyed sites around Cozumel.

**ESM 3.**
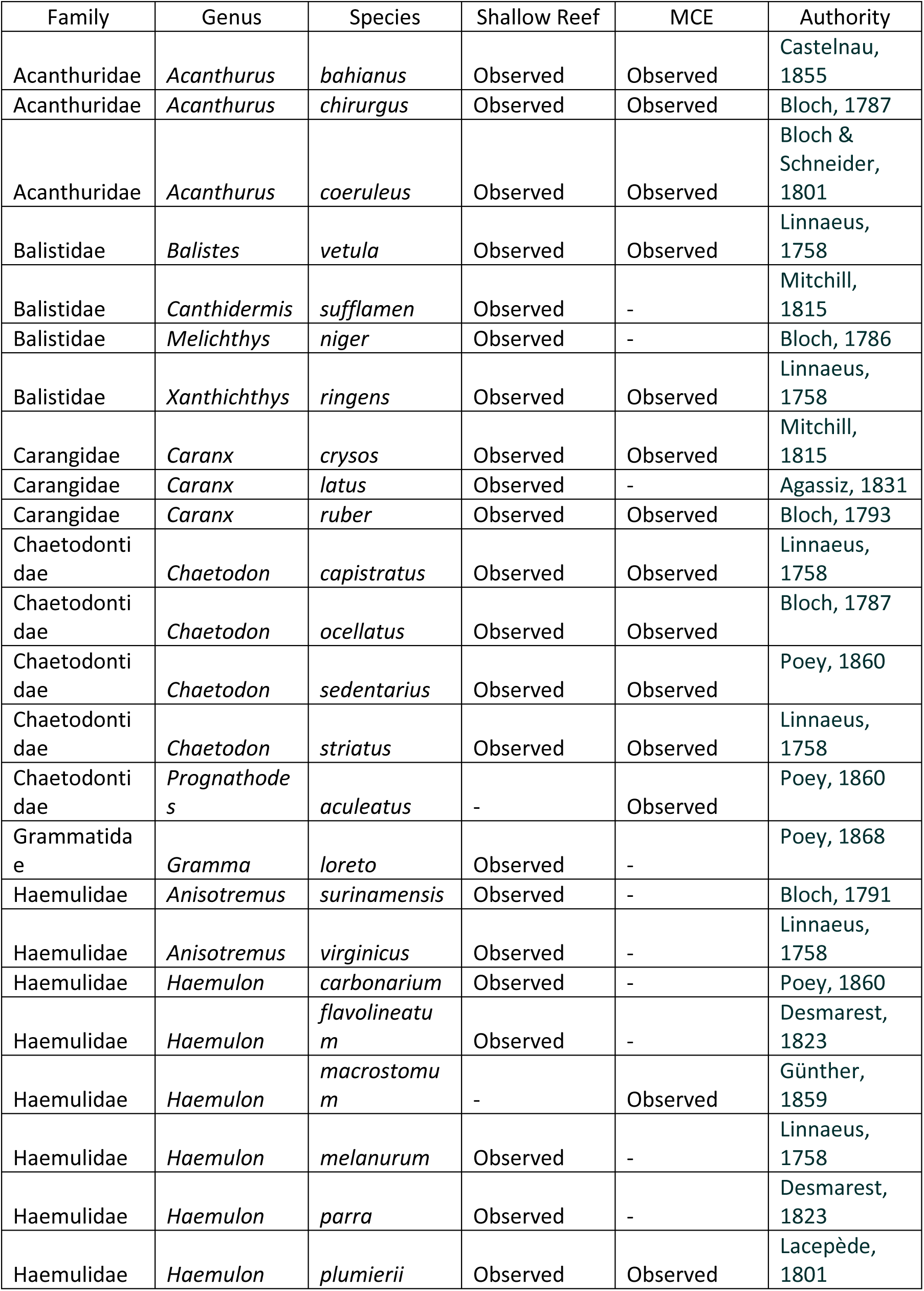
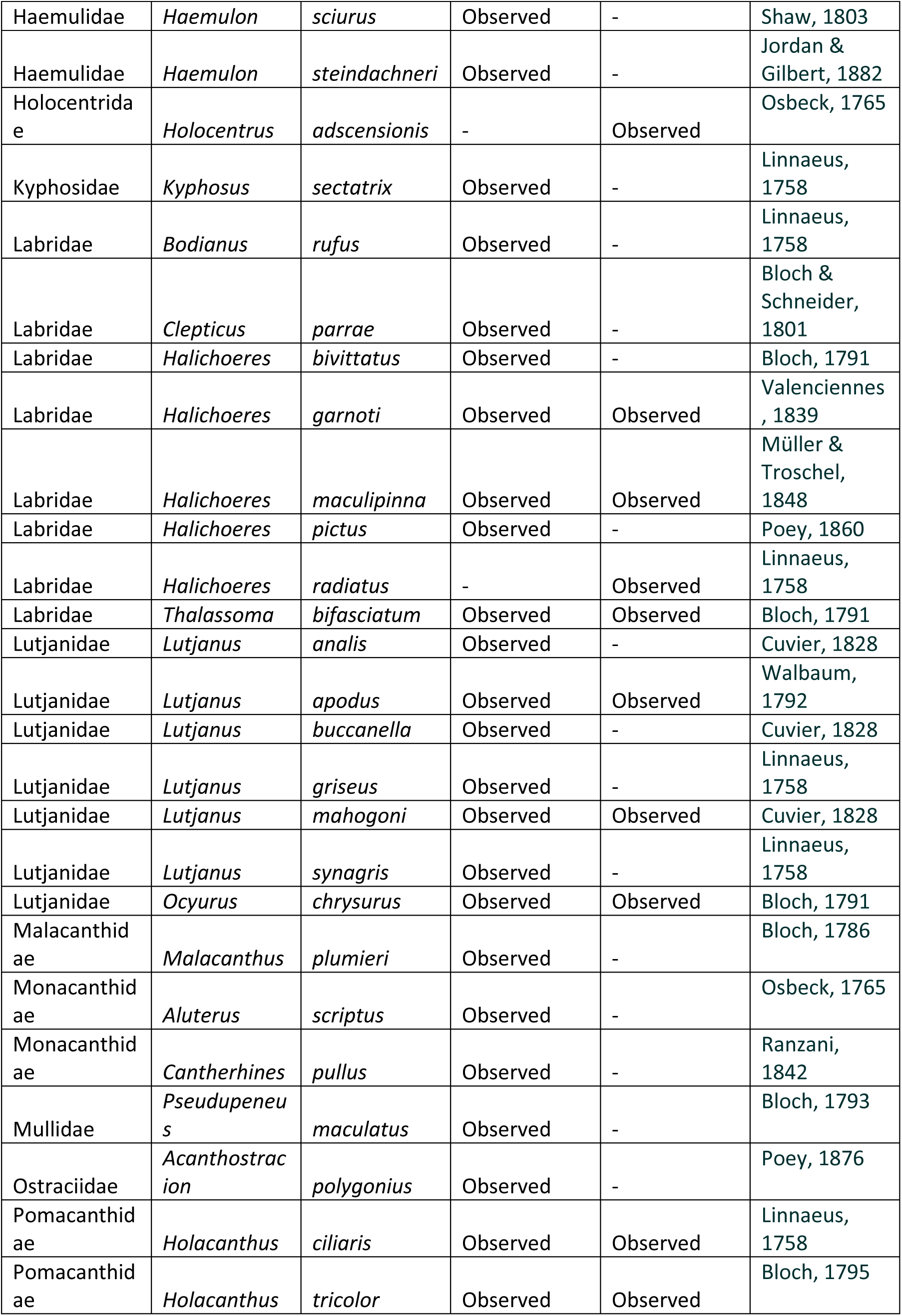
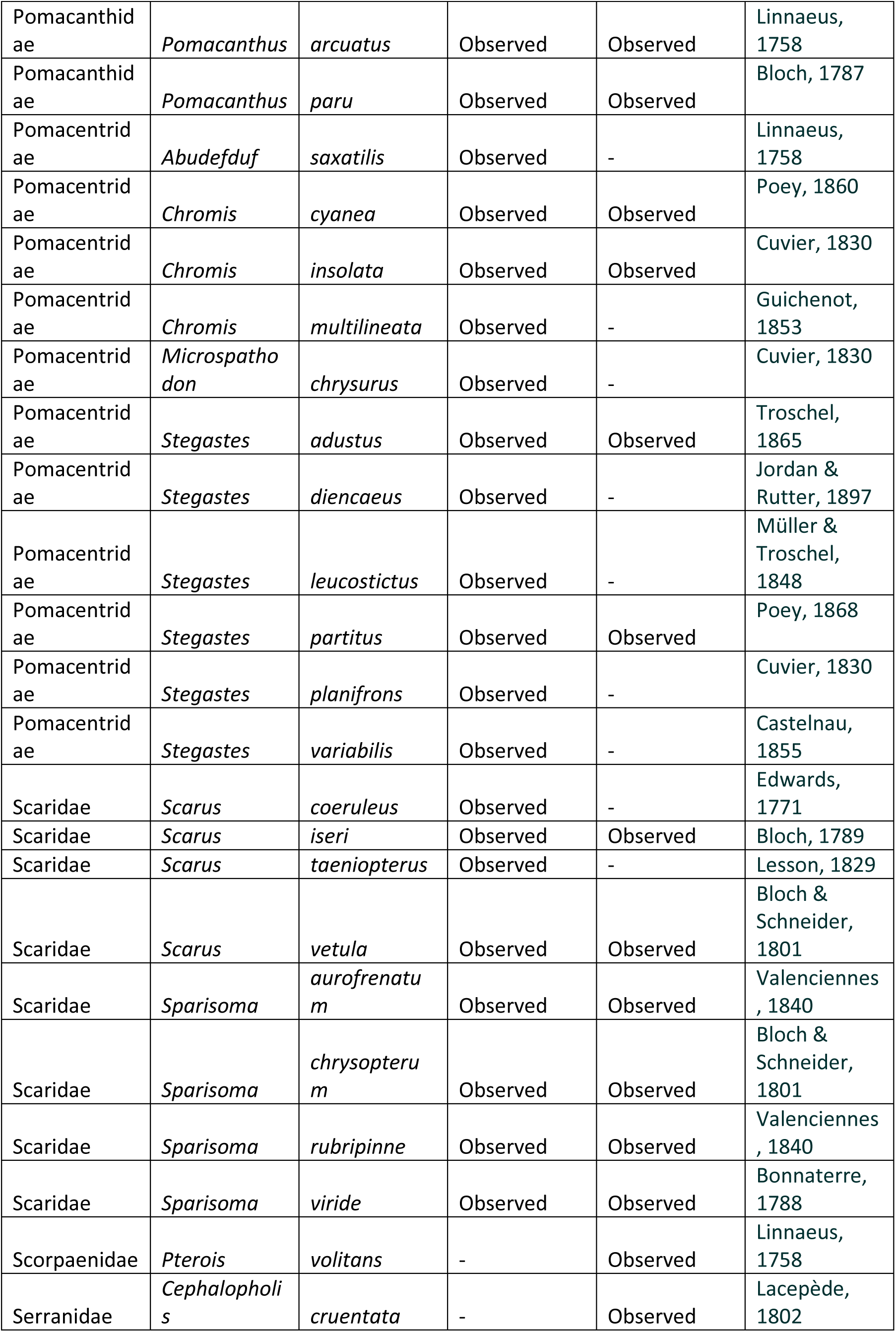
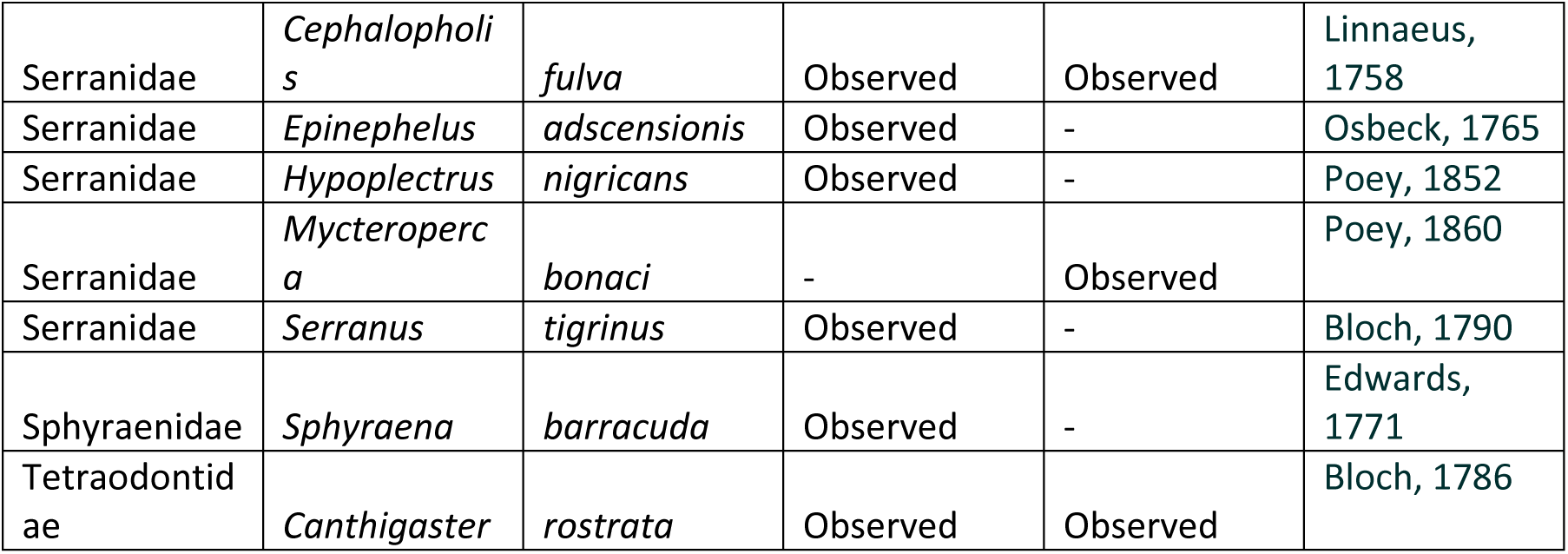
Fish species observed on shallow reefs (15 m) and MCEs (55 m) at surveyed sites around Cozumel.

